# Squamous differentiation portends poor prognosis in low and intermediate-risk endometrioid endometrial cancer

**DOI:** 10.1101/698316

**Authors:** Diocésio Alves Pinto de Andrade, Vinicius Duval da Silva, Graziela de Macedo Matsushita, Marcos Alves de Lima, Marcelo de Andrade Vieira, Carlos Eduardo Mattos Cunha Andrade, Ronaldo Luís Schmidt, Rui Manuel Reis, Ricardo dos Reis

## Abstract

**Background:** Endometrial cancer presents well-defined risk factors (myometrial invasion, histological subtype, tumor grade, lymphovascular space invasion (LVSI)). Some low and intermediate-risk endometrioid endometrial cancer patients exhibited unexpected outcomes. The aim of this study was to investigate other clinical-pathological factors that might influence the recurrence rates of patients diagnosed with low and intermediate-risk endometrioid endometrial cancer.

**Methods:** A case-control study from a cohort retrospective of 196 patients diagnosed with low and intermediate-risk endometrioid endometrial cancer at a single institution between 2009 and 2014 was conducted. Medical records were reviewed to compare clinical (race, smoking, menopause age, body mass index) and pathological (histological characteristics (endometrioid *vs* endometrioid with squamous differentiation), tumor differentiation grade, tumor location, endocervical invasion, LVSI) features of patients with recurrence (case) and without recurrence (control) of disease. Three controls for each case were matched for age and staging.

**Results:** Twenty-one patients with recurrence were found (10.7%), of which 14 were stage IA, and 7 were stage IB. In accordance, 63 patients without recurrence were selected as controls. There were no significant differences in any clinical characteristics between cases and controls. Among pathological variables, presence of squamous differentiation (28.6% *vs.* 4.8%, p=0.007), tumor differentiation grade 2 or 3 (57.1% *vs.* 30.2%, p=0.037) and presence of endocervical invasion (28.6% *vs.* 12.7%, p=0.103) were associated with disease recurrence on univariate analysis. On multivariable analysis, only squamous differentiation was a significant risk factor for recurrence (p=0.031).

**Conclusion:** Our data suggest that squamous differentiation may be an adverse prognostic factor in patients with low and intermediate-risk endometrioid endometrial cancer, that showed a 5.6-fold increased risk for recurrence.

## Introduction

Endometrial cancer is the most prevalent gynecological neoplasia in women in the US, accounting for more than 63,000 cases/year and with a lethality rate close to 18%.^1^ In Brazil, this tumor represents the second cause of gynecological cancer due to a high incidence of tumors of the cervix.^2^ Despite knowledge advances related to genetic alterations of this neoplasia in the last years, classification of endometrial cancer into type I (endometrioid) or type II (serous or clear cell) continues to be used in clinical practice, mainly to evaluate risk factors in tumor development.^3^

As in other solid tumors, staging of endometrial cancer is important to define surgical extension, ranging from hysterectomy with bilateral salpingo-oophorectomy even need pelvic and/or para-aortic lymphadenectomy.^4^ Risk stratification in stage I tumors aims to assess the risk of lymph node involvement, the recurrence pattern, patient’s prognosis and the best adjuvant treatment to be performed.^5^ Beyond myometrial tumor invasion depth, other clinical-pathological factors were evaluated: age; histological subtype; tumor differentiation grade and lymphovascular space invasion (LVSI).^6, 7^ Beside these features, other immunohistochemistry markers, such as L1-cell adhesion molecule (L1CAM) and p53 are also associated with patient outcome for stage I endometrial cancer, but not yet incorporated in the current classifications.^8, 9^

Endometrial adenocarcinoma with squamous differentiation terminology was defined by Zaino and Kurman in 1988 to replace two previously used nomenclature for uterus neoplasms: adenoacanthoma and adenosquamous carcinoma.^10^ Squamous differentiation consists of sheets of cells with intercellular bridges and prominent cell membranes with or without keratinization.^11^ It is present in about 13-25% of endometrial adenocarcinomas.^10, 12^ The finding of squamous differentiation in the anatomopathological examination remains controversial as a risk factor for recurrence in patients with early stage endometrial cancer.^13, 14^ The aim of this study was to evaluate clinical-pathological features that influenced the recurrence of patients diagnosed with low and intermediate risk endometrial cancer according to ESMO (European Society for Medical Oncology) criteria.^5^

## Patients and Methods

A case-control study nested in a retrospective cohort of 196 patients diagnosed with low and intermediate risk endometrial cancer undergoing surgery at Barretos Cancer Hospital between January 2009 and December 2014 was conducted. This study was conducted in accordance with the principles of the Declaration of Helsinki, and was previously approved by the Ethical Review Board from Barretos Cancer Hospital in March 2017 (Reference 1.942.488). Cases were defined as patients who presented systemic or locoregional recurrence at any time of their follow up. We defined three controls for each recurrence case matched age (± 1 year) and FIGO (International Federation of Gynecology and Obstetrics) staging (IA and IB).

According to ESMO criteria^5^, low-risk endometrial cancer is defined as endometrioid adenocarcinoma stage IA grade 1 or grade 2; intermediate-risk endometrial cancer is defined as endometrioid adenocarcinoma stage IA grade 3 or endometrioid adenocarcinoma stage IB grade 1 or grade 2. Three or more of the following four criteria need to be present to define squamous differentiation: sheet-like growth without glands or palisading, sharp cells margins, eosinophilic and thick of glassy cytoplasm, and decreased nuclear-to-cytoplasmic ratio compared with foci elsewhere in the same tumor.^11^ The amount of squamous differentiation can vary, and in a well-sampled carcinoma, the squamous differentiation should comprise at least 10% of the neoplasia. The degree of nuclear atypia, if present, generally reflects that of the glandular cells.^15^

Clinical-pathological data were reviewed from medical records. The diagnoses of low and intermediate risk endometrial cancer were confirmed by surgical histopathologic report. Patients who did not perform definitive surgical treatment at the institution (for example, patients who underwent surgery at their region of origin and who were referred to a tertiary hospital only for adjuvant treatment) were excluded.

The following clinical-pathological criteria were evaluated: ECOG (Eastern Cooperative Oncology Group) scale of performance status (0-1 vs 2); race/ethnicity (white *vs* non-white); body mass index (BMI); hormonal status (menopause *vs* menacme); number of pregnancies; smoking (yes *vs* no); tumor differentiation grade (1, 2 or 3); histological characteristics (endometrioid *vs* endometrioid with squamous differentiation); tumor size; tumor location (uterine corpus *vs* lower uterine segment); endocervical invasion (yes *vs* no) and LVSI (yes *vs* no).

### Statistical Analysis

Both the data collected and analyses were performed using IBM Statistical Package for the Social Sciences (SPSS) database version 21.0 (SPSS, Chicago, IL). Descriptive statistical analysis used median, maximum and minimum value for quantitative variables and percentage for qualitative variables. Once the above variables were defined, univariate analysis was performed using Mann-Whitney’s U-test or Fisher’s exact test. Parameters with *P* < 0.2 in univariate analyses were entered into the logistic regression analysis. Backward stepwise logistic regression models were constructed. The comparisons were considered statistically significant at *P* < 0.05. Study data were collected and managed using REDCap (Research Electronic Data Capture) electronic data capture tools hosted at Barretos Cancer Hospital.^16^

## Results

Of the 196 endometrial cancer patients described in this retrospective cohort, 21 patients (10.7%) presented recurrence during their evolution (cases), of which 2/3 were stage IA and 1/3 were stage IB, and 63 patients without recurrence were selected as controls (Table 1). The median age of both groups was 64 years and both groups also exhibit similar fraction of IA staging. Moreover, the patient population was obese (median BMI above 30), white and was non-smoker (Table 1). Almost all patients were already in menopause (11.2% of patients controls were still in menacme).

**Table 1.** Univariate analysis of predictive recurrence for low and intermediate-risk endometrioid endometrial cancer.

|  |  | Case (n=21) | Control (n=63) | P-value |
| --- | --- | --- | --- | --- |
| Age (median)<sup>a</sup> |  | 64 (46-77) | 64 (46-78) | 0.873 |
| FIGO staging (%)<sup>b</sup> | IA | 14 (66.7) | 42 (66.7) | >0.99 |
|  | IB | 7 (33.3) | 21 (33.3) |  |
| ECOG Performance Status (%)<sup>b</sup> | 0-1 | 20 (95.2) | 61 (96.8) | >0.99 |
|  | 2 | 1 (4.8) | 2 (3.2) |  |
| Race/Ethnicity (%)<sup>b</sup> | White | 18 (85.7) | 45 (71.4) | 0.251 |
|  | Non-white | 3 (14.3) | 18 (28.6) |  |
| BMI (median)<sup>a</sup> |  | 31.64 (19.78-48.62) | 32.65 (21.93-52.71) | 0.339 |
| Smoking history<sup>b</sup> | Yes | 2 (9.5) | 4 (6.3) | 0.637 |
|  | No | 19 (90.5) | 59 (93.7) |  |
| Menopause (%)<sup>b</sup> | Yes | 21 | 56 (88.8) | 0.184 |
|  | No | 0 | 7 (11.2) |  |
| Number of pregnancies (median)<sup>a</sup> |  | 4 (1-7) | 4 (1-20) | 0.725 |
| Tumor differentiation grade<sup>b</sup> | Grade 1 | 9 (42.9) | 44 (69.8) | 0.037 |
|  | Grade 2 or 3 | 12 (57.1) | 19 (30.2) |  |
| Histological subtype (%)<sup>b</sup> | Endometrioid | 15 (71.4) | 60 (95.2) | 0.007 |
|  | Endometrioid with squamous differentiation | 6 (28.6) | 3 (4.8) |  |
| Tumor size (median – cm)<sup>a</sup> |  | 4.0 (16.0-115.0) | 4.0 (1.0-105.0) | 0.597 |
| Tumor localization<sup>b</sup> | Uterine corpus | 14 (66.7) | 47 (74.6) | 0.574 |
|  | Lower uterine segment | 7 (33.3) | 16 (25.4) |  |
| Endocervical invasion (%)<sup>b</sup> | Yes | 6 (28.6) | 8 (12.7) | 0.103 |
|  | No | 15 (71.4) | 55 (87.3) |  |
| LVSI (%)<sup>b</sup> | Yes | 5 (23.8) | 9 (14.3) | 0.324 |
|  | No | 16 (76.2) | 54 (85.7) |  |

BMI – body mass index; ECOG – Eastern Cooperative Oncology Group; FIGO – International Federation of Gynecology and Obstetrics; LVSI – lymphovascular space invasion. a- Mann-Whitney test; b- Fisher's exact test

Squamous differentiation appears as solid areas in the middle of glandular tissue. These areas, although solid, can not be considered as such for grading purpose (Figure 1a and 1b). A specific immunohistochemical marker used to evaluate squamous lineage is p63, as shown in the inset (Figure 1c).^17^

**Figure 1.**
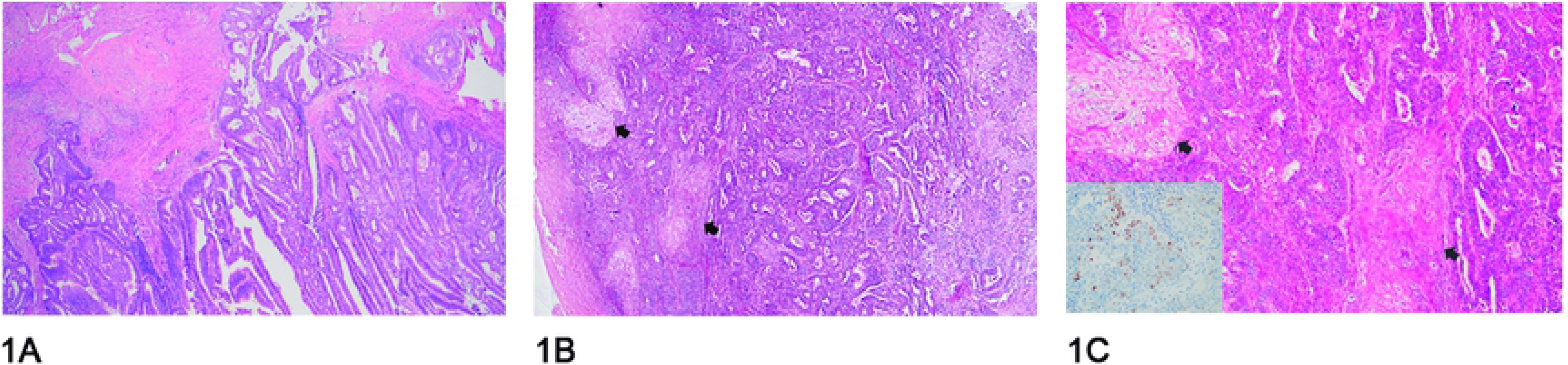
Figure 1A depicts an endometrioid adenocarcinoma without squamous transformation, 1B shows a case with squamous transformation areas highlighted with arrows and 1C highlights the squamous transformation areas at a higher magnification (arrows). The inset presents nuclear p63 positivity, a protien antibody used to demonstrate squamous differentiation by immunohistochemistry in a squamous transformation area.

There were no significant differences in race/ethnicity, ECOG performance status, number of pregnancies, smoking history, tumor size, tumor localization and LVSI between the group of patients with recurrence (cases) and patients without recurrence (controls) (Table 1).

In the univariate analysis, four parameters with *P* < 0.2 were chosen for the multivariate logistic regression analysis: hormonal status (menopause), tumor differentiation grade, histological characteristics and endocervical invasion (Table 1). The variable menopause had to be withdrawn from this model since one of its categories did not present participants (no menopause in case group), resulting in a no data conversion to the odds ratio value. Using backward stepwise logistic regression technique, a new model were constructed with three parameters: histological subtype with squamous differentiation (28.6% *vs.* 4.8%, p=0.007), tumor differentiation grade 2 or 3 (57.1% *vs.* 30.2%, p=0.037) and presence of endocervical invasion (28.6% *vs.* 12.7%, p=0.103) (Table 1).

In multivariate analysis, only histological subtype (endometrioid vs endometrioid with squamous differentiation) was associated with recurrence (p=0.031) (Table 2). Women who presented squamous differentiation associated with classic endometrioid subtype had a 5.6-fold increased risk for recurrence when compared to the group that does not show this histological finding (Table 2).

**Table 2.** Multivariate analysis of predictive recurrence for low and intermediate-risk endometrioid endometrial cancer.

|  |  | **Odds Ratio (IC – 95%)** | **P-value** |
| --- | --- | --- | --- |
| Tumour differentiation grade | Grade 1 | 1 | 0.080 |
|  | Grade 2 or 3 | 2.66 (0.89–7.96) |  |
| Tumour type | Endometrioid | 1 | 0.031 |
|  | Endometrioid with squamous differentiation | 5.65 (1.17–27.17) |  |
| Endocervical invasion | No | 1 | 0.168 |
|  | Yes | 2.55 (0.67–9.66) |  |

Constant= -1.939 ( $P = 0.0001$ )

## Discussion

This case-control study of low and intermediate risk endometrial cancer demonstrated that patients with endometrioid squamous differentiation subtype had a greater chance of recurrence when compared to patients with typical endometrioid histological subtype. This finding in the anatomopathological examination remains controversial as a risk factor for recurrence as published in the international literature (Table 3).

**Table 3.** Summary of squamous differentiation endometrioid endometrial cancer studies to predict recurrence.

| References | Year | Country | N | Study design | Risk for recurrence |
| --- | --- | --- | --- | --- | --- |
| This study | 2019 | Brazil | 84 | Case-control | Yes |
| Misirlioglu *et al.*<sup>13</sup> | 2012 | Turkey | 223 | Case-control | Yes |
| Jiang *et al.*<sup>21</sup> | 2017 | China | 630 | Retrospective cohort | Yes |
| Zaino *et al.*<sup>14</sup> | 1991 | USA | 631 | Prospective cohort | No |
| Sturgeon *et al.*<sup>22</sup> | 1998 | USA | 648 | Case-control | No |
| Lax *et al.*<sup>23</sup> | 1998 | USA | 77 | Case series | Variable |
| Abeler *et al.*<sup>12</sup> | 1992 | Norway | 255 | Retrospective cohort | Variable |

FIGO staging classifies endometrial cancer grade into three groups: grade 1 tumors are those in which less than 5% of the neoplasm is arranged as solid growth; grade 2 tumors are those in which 5% to 50% of the neoplasms are arranged in solid sheets, and grade 3 tumors are those in which greater than 50% of the neoplasm form solid masses.^18^ The current FIGO grade system, primarily based on the relative proportion of solid and glandular areas also considers nuclear atypia, and grading is increased by one if more than 50% severe nuclear atypia (grade 3 nuclei) is found in the neoplastic glands.^19^ Currently, squamous differentiation does not enter into this classification, although it can mimic solid tumors areas. It can be found in all forms of endometrial hyperplasia, being more common in atypical endometrial proliferation.^15^ The squamous and glandular components have the same PTEN mutations, which indicates that they are clonally related.^20^

Some studies showed that squamous differentiation is a risk factor for endometrial cancer recurrence.^13, 21^ A retrospective study of 223 patients with early-stage endometrial cancer, carried out by Misirlioglu *et al.*, similar with our study, regarding methodological structure, showed squamous differentiation as a risk factor for recurrence in early-stage endometrial cancer.^13^ The authors reported 10.31% of recurrence (23 cases), very similar to that found in our study. Several risk factors were considered positive to increase the chance of recurrence (age, depth of myometrial tumor invasion, tumor differentiation grade, lymphovascular space invasion, tumor localization, tumor size), including squamous differentiation as in our results.^13^ Another retrospective cohort with 630 patients with stage I endometrioid endometrial cancer conducted by Jiang *et al.* evaluated possible risk factors for metastasis in this tumor. Beyond traditional factors such as tumor size and depth of myometrial invasion, squamous differentiation was also an independent risk factor for the development of pulmonary metastasis.^21^

On the other hand, there are some studies showing that squamous differentiation does not pose a worse prognosis. A large study (n=631) conducted by Gynecologic Oncology Group (GOG) in the late 1970s and early 1980s, evaluated the prognosis role of the patients with or without histological squamous differentiation.^14^ Five-years overall survival was 90% for patients with squamous differentiation *versus* 82% for patients without this differentiation with statistical significance.^14^ A case-control study with 640 patients carried out by Sturgeon *et al.* showed that squamous differentiation is not a poor prognostic factor for patients diagnosed with endometrioid endometrial cancer.^22^

On account of conflicting results for defining prognosis of tumors; it may be necessary to classify squamous differentiation component in the low or high degree. An immunohistochemistry study of 77 patients evaluated estrogen (ER) receptor, progesterone (PR) receptor, p53 and Ki-67, reported that tumors with high-grade squamous differentiation (lack of expression of ER and PR; high Ki-67 index and p53 expression) have a worse outcome.^23^ This controversy about the prognosis of recurrence in endometrial cancer with squamous differentiation may be related to subgroups of its classification. Abeler *et al.* published a cohort with 1985 cases with endometrioid endometrial carcinoma, of which 255 presented squamous differentiation.^12^ In this study, the authors divided tumors with squamous differentiation into two groups formerly used: adenoacanthoma (for cytologically well differentiated squamous differentiation) and adenosquamous carcinoma (for poorly differentiated squamous differentiation). Five-year overall survival for all patients was 83.5%. Adenoacanthoma subgroup had 91.2% five-year overall survival and adenosquamous subgroup had 64.9%, showing different prognosis.^12^

Molecular analysis with the aim to discover a biomarker that correlates with squamous differentiation in endometrial cancer is even more unclear. Cdx2 is an important gene transcription factor in the carcinogenesis of colorectal cancer.^24^ The expression of this biomarker can be present in up to 27% of endometrial cancer but it is never seen in the normal epithelium.^25^ Wani *et al.* evaluated Cdx2 expression in endometrial cancer with or without squamous differentiation and the expression of the biomarker was more prevalent in patients with this differentiation.^25^ Another biomarker that may be related to squamous differentiation in endometrial cancer is p16, a tumor suppression protein generally expressed in tumors caused by the human papillomavirus (HPV).^26, 27^

The strengths of our study include the fact that all patients were treated at the oncoginecology department from a tertiary cancer hospital where protocols are followed closely. The pathology department is also divided into subspecialties, surgical specimens description, sampling, and reporting are standardized, resulting in high reproducibility of the pathology reports. Furthermore, the methodology chosen was a well-matched case-control study by age and stage, without differences between groups.

The limitations of the present study is its retrospective nature, associated with the number of recurrent cases found (10.71%), despite agreeing with data from literature since it is low and intermediate risk stage I tumors.^13^ Creasman *et al.* reported a relapse-free survival at five years in stage I surgical patients of 92.3%.^28^ Tumor differentiation grade and endocervical invasion were not statistically significant in the multivariate analysis model, probably due to this limitation. Other barriers of this study were to have been carried out in a single institution with possible referral bias, and it did not have any immunohistochemical data.

In conclusion, this case-control study provides evidence that squamous differentiation in low and intermediate risk endometrial cancer had a 5.6-fold increased risk for recurrence. This finding demonstrates that more detailed histopathological information could contribute to the analysis of prognosis for the patients.

## Disclosures

The authors do not have any conflicts of interest to disclose.

